# Comparing methanol-glucose and dimethyl-sulfoxide based extender for milt cryopreservation of brown trout (*Salmo trutta*)

**DOI:** 10.1101/289736

**Authors:** David Nusbaumer, Lucas Marques da Cunha, Claus Wedekind

**Affiliations:** Department of Ecology and Evolution, Biophore, University of Lausanne, Lausanne, Switzerland

**Keywords:** Sperm, cryopreservation, methanol, glucose, brown trout

## Abstract

The potential importance of sperm cryopreservation for aquaculture and conservation management seems still undervalued, probably because the available protocols often lead to reduced fertilization success. We experimentally compared the effectiveness of two different freezing extenders for cryopreservation of brown trout (*Salmo trutta)* semen, controlling for possible male and female effects. The methanol-glucose based extender that we tested was significantly more effective than a common dimethyl-sulfoxide based extender (a commercial cryopreservation kit). We then studied the effectiveness of the methanol-glucose based extender at different sperm-egg ratios and found no significant differences in fertilization ability of fresh and cryopreserved milt at a sperm-egg ratio of at least 110,000:1. We conclude that brown trout sperm cryopreserved with this extender can be used even at low sperm-egg ratios without significant effects on fertilization rates.

## 1. Introduction

Minimizing damage to sperm in cryopreservation of fish semen has been a target for decades of research and is important in aquaculture and in conservation biology. Effective cryopreservation can ensure availability of gametes when there is desynchronization between male and female breeders, and can play an important role in the conservation of rare breeds or in the genetic improvement of a cultured population [1]. Moreover, cryopreservation with minimal damage to sperm would be the basis of sperm banks that preserve the genetic resources of a threatened population, such as the National Animal Germplasm Program (NAGP, http://ars.usda.gov/research/projects/projects.htm?accn_no=423549). Sperm can then be used to support restocking programs while minimizing disturbance at the spawning ground or to reduce inbreeding depression or hybridization of a population [2,3].

Since the first attempts, much progress has been made and effective protocols now exist for a variety of fishes, mainly freshwater fishes [1]. Salmonids have been a focus of these research efforts due to their commercial and cultural importance [1]. The first successful cryopreservation was achieved using glycerol as a cryoprotectant [4], but it was quickly replaced by dimethyl sulfoxide (DMSO). Most of the current protocols still use DMSO as permeable cryoprotectant, and DMSO is still considered as a suitable candidate for the development of new protocols. For instance, DMSO was used lately in the development of a protocol for the endangered Mediterranean brown trout *Salmo trutta macrostigma* [5]. However, methanol was suggested as an alternative to DMSO and tested in three salmonids including brown trout [6]. Permeable cryoprotectants are most often associated with complex saline solutions and non-permeable cryoprotectants, such as egg yolk. Recently, a very simple extender consisting only of 9% methanol and 0.15M glucose was shown to be effective in rainbow trout (*Oncorhynchus mykiss*) and in brown trout [7,8].

Here we compare the effectiveness of this methanol-glucose based extender to a DMSO-based extender on brown trout while controlling for potentially confounding parental or population effects on fertilization success. The DMSO-based extender we use here is a commercial product that use DMSO as permeable cryoprotectant, egg yolk derived lipids as non-permeable cryoprotectant, and a saline solution. We ran two experiments. The first one aimed to assess which of the two candidate extenders would produce the highest fertilization success. In the second experiment, we tested the post-thaw fertilizing ability of sperm at various dilutions.

## 2. Methods

### 2.1 Collection of gametes

Wild males and female brown trout were caught by electrofishing in different tributaries of the Aare river and kept in the facilities of the *Fischereistützpunkt* Reutigen until collection of the gametes (either on 18.11.2015 or on 2.12.2015). Milt was stripped drop by drop into 145×20mm Petri dishes. Care was taken to avoid drops mixing. Milt from drops that did not seem to be contaminated by urine or feces was collected with a pipette and stored (< 1h) on ice in a 2ml micro tube (Sarstedt, Germany) until preparations for cryopreservation started. Eggs were stripped into plastic containers from which 8 eggs per female were separated in 60 x 15mm Petri dish (Greiner bio-one, Germany) and stored at ambient temperature (4-7°C) for < 30 min until fertilization.

### 2.2 Sperm cryopreservation

Two freezing extenders were used: i) Cryofish (IMV Technologies, France) and ii) methanol 10% + glucose 0.15M. The first was prepared mixing the following kit solutions: 8 volumes of Freezesol (saline solution) + 1 volume of DMSO + 1 volume of Freezlip (lipids solution meant to replace egg yolk). The second extender was prepared by adding 20 mL of methanol (VWR chemicals, Switzerland) and 5.945 g of D-glucose monohydrate (Fisher chemicals, Switzerland) to 180 mL of ultrapure water [7]. Both extenders were kept on ice before use. Microtubes were prepared with 300 µL Cryofish (following manufacturer instructions) or 500 µL Methanol extender (following recommendations of [7]).

For the first experiment, two 100 µL samples of milt per male were added to one of the two extenders, respectively 300 µL of Cryofish or 500µl of methanol-glucose, and vortexed for 5 seconds. The samples on Cryofish were then immediately processed further (following the manufacturer’s instructions and because DMSO is toxic to the sperms [9]) while samples in the methanol-glucose based extender were given a 15-minutes equilibration time on ice before freezing. In the second experiment, only the latter extender was used at a 1:5 ratio (100 µl milt in 500 µl extender).

Two 66.5 mm CTE straws (MTG Technologies, Germany) were used per milt sample and extender. They were filled with 200 µL of a mix each, sealed at both end with a straw sealer (MTG Technologies, Germany), and kept on ice until freezing. For freezing, straws were first placed for 15 minutes on a floating rack within the liquid nitrogen tank, about 1.5 cm above the surface of the liquid, before they were plunged into liquid nitrogen. Micro tubes containing fresh sperm were kept on ice until fertilization (< 45 min).

### 2.3 Fertilization

In the first experiment, fresh and cryopreserved milt of five males were used to fertilize eggs of 2 females each. This breeding design allowed to fertilize 80 eggs per treatment while controlling for parental effects (Figure 1). Straws were individually removed from the liquid nitrogen, plunged for 30 seconds in water at 25°C and put on ice for 1 minute. Then, the content of the straw was dropped into a Petri dish with the respective egg sample, not mixing milt and eggs yet. In parallel, 33 µL of fresh milt of the same male was similarly placed around the other egg sample. We used 33 µL as this is corresponds to the absolute volume of milt contained in one straw filled with the methanol based extender. Fresh sperm and frozen-thawed sperm were then activated and mixed with their respective egg sample by adding 4 ml of Actifish solution (IMV Technologies, France) to each Petri dish (i.e. 500 µl solution per egg). The Petri dishes were then gently moved to support the mixing of the gametes. After 5 minutes, 5 mL of standardized water [10] was added to each Petri dish and the eggs were left undisturbed for 2 hours to allow hardening.

**Figure 1.**
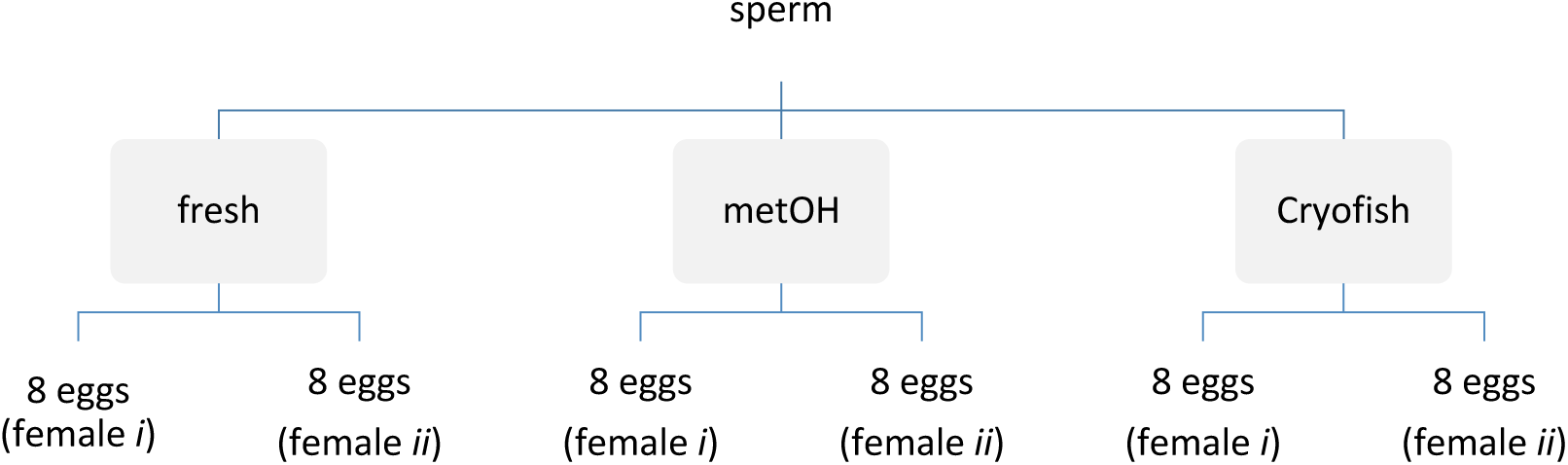
Design of one breeding block (n=5) in the first experiment.

In the second experiment, the same thawing procedure as described above was followed before straws were emptied into 2 mL microtubes on ice (2 straws per tube). Three serial dilutions of 400 µL were then made following a 10-fold decrease (100%, 10% and 1%) in the extender for the frozen-thawed milt and in Storfish (IMV Technologies, France) for fresh milt. The 100% dilution referred to milt already diluted at 1:5 in the extender. Therefore, concentration of the control (fresh milt) was adjusted accordingly. This led to final dilution of 16.5%, 1.65% and 0.165%, implying an absolute volume of milt of 66, 6.6 and 0.6 µL of milt in extender or Storfish, respectively. We used 200 µL of each dilution to fertilize the eggs following the same procedure as described above. Thus, every batch of eggs was fertilized with an absolute volume of 33, 3.3 or 0.33 µL of either fresh or frozen thawed milt. We tested fresh and cryopreserved milt of in total 4 males with eggs of 2 females each. This breeding design allowed us to fertilized 192 eggs per treatment and 64 eggs per dilution within treatment, while controlling for parental effects (Figure 2). Sperm activation was done as in the first experiment.

**Figure 2.**
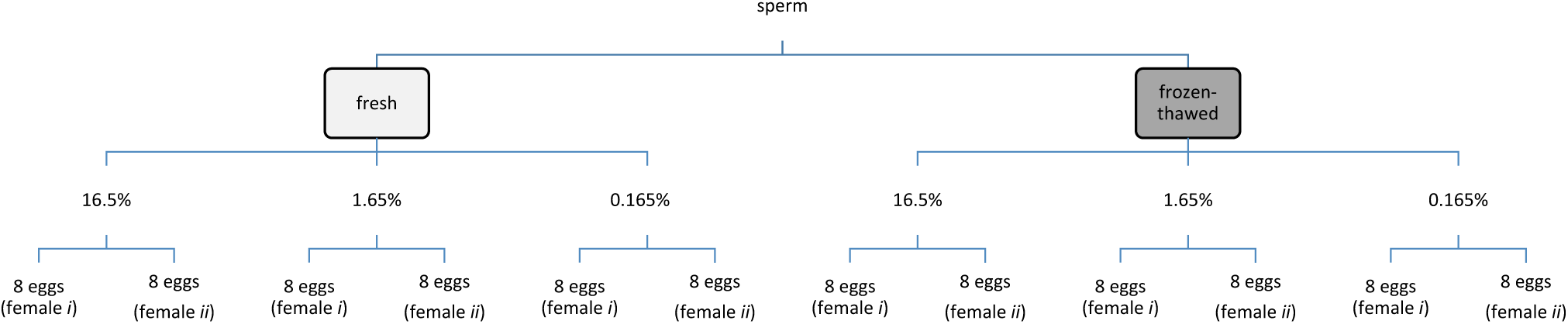
Design of one breeding block (n=4) in the second experiment.

### 2.4 Transportation to the laboratory and distribution of the eggs

After hardening, eggs were transferred to 50 ml conical (Greiner Bio-one, Germany) with approximately 30ml standardized water and transported on ice to the laboratory (2 hours). There, each tube was emptied in a plastic tea strainer and the eggs were placed in a 145 mm Petri dish filled with autoclaved standardized water. Eggs were then distributed singly to wells of 24-well plates filled with 1.8ml autoclaved standardized water per well. Plates were incubated at 7°C in a climate chamber at a 12-hours light cycle. After 13 days, the fertilization success was assessed with a light table. Eggs were considered fertilized if the spinal cord of the embryo was visible. Eggs were called unfertilized if no embryo was visible at that time point.

### 2.5 Sperm concentration

We used the CASA software (Qualisperm®, Biophos SA, Switzerland) to assess sperm concentration of fresh milt to calculate the actual amount of sperm cells in the different dilution of the density experiment. Therefore, 20 µL of milt were added to 180 µL of Storfish in a 2 ml test tube, kept on ice, and transported to the lab. There, milt was diluted again to 1:500 with standardized water. From this, 2 µL were transferred in a 4-well chamber slide (Leja, Netherlands) on a cooling stage set at 6.5°C. Sperm was observed at 20x magnification and with phase contrast. Concentration was given by the program in mio/ml.

### 2.6 Statistics

Fertilization success was analyzed in generalized linear mixed effect models with the lme4 package [11] in Rstudio [12]. For the first experiment, treatment (type of extender) was entered as a fixed factor in the model, while male and female identities were entered as random factors. For the second experiment, treatment (fresh vs. frozen-thawed) and dilution (16.5, 1.65 and 0.165 %) were entered as fixed factors, while male and female identity were again entered as random factor as well as their interactions with the fixed effects. To test the significance of an effect, a model including or lacking the term of interest was compared to the reference model. The goodness of fit of the different models is given by the logarithm of the approximated likelihood and by the Akaike’s information criterion. To test if models differ in their goodness of fit, the models were compared with likelihood ratio tests (LRT). For treatments that had more than two levels, we also ran a multiple comparison of means on the reference model using Tukey method with the multcomp package [13] in Rstudio.

## 3. Results

### 3.1 First experiment

We found treatment and female identity to significantly affect the fertilization rates (Table 1). Sperm cryopreserved with Cryofish led to reduced fertilization rate when compared to fresh sperm (z = -2.5; p = 0.03) and to sperm cryopreserved with MetOH (z = 2.9; p = 0.009). However, the latter did not lead to reduced fertilization success when compared to fresh sperm (z = 0.4; p = 0.89) (Figure 3).

**Table 1.** Likelihood ratio tests on mixed-effects model regressions on fertilization success. Models including or lacking the term of interest were compared to reference models in bold to determine the significance of the effect tested.

| Model terms | Effect tested | AIC | d.f. | $X^2$ | *P* |
| --- | --- | --- | --- | --- | --- |
| **t+b+f** |  | 226 | 5 |  |  |
| b+f | t | 233 | 3 | 10.4 | **0.005** |
| t+f | m | 224 | 4 | 0 | 1 |
| t+b | f | 260 | 4 | 36 | **<0.001** |

Fixed effects: **t**, treatment; Random effects: **b**, male; **f**, female. $P<0.05$ are shown in bold

**Figure 3.**
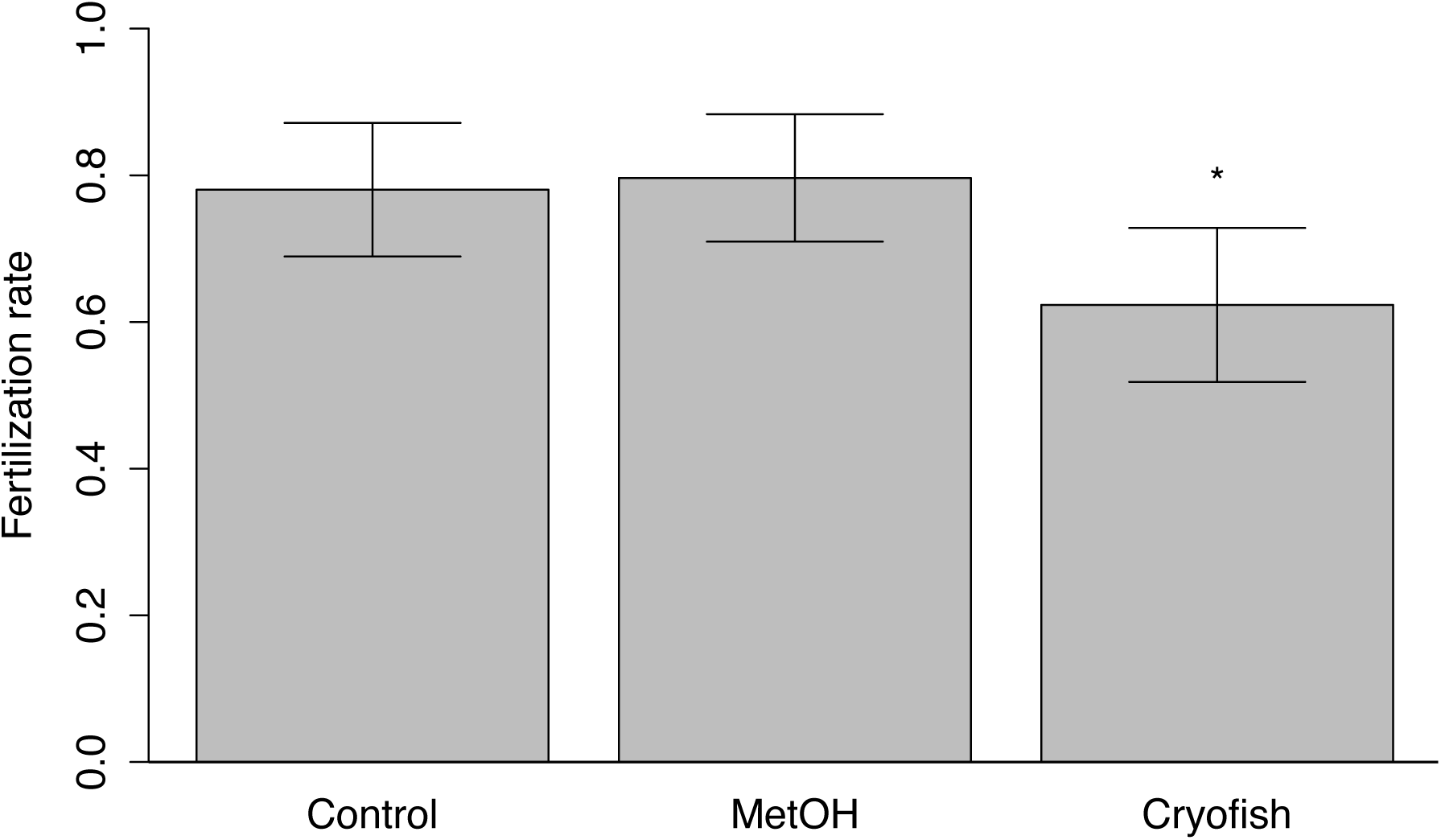
Mean fertilization success at 13 days post fertilization (dpf) in the first experiment. Error bars indicate 95% confidence interval.

### 3.2 Second experiment

Both treatment (cryopreservation) and dilution had a significant effect on fertilization (Table 2). Significant differences were found only between fresh-thawed sperm diluted at 0.165% and all the other groups: against fresh-thawed 16.5% (z = 5.3; p < 0.001), against fresh-thawed 1.65% (z = 4.5; p < 0.001), against control 16.5% (z = 5.0; p < 0.001), against control 1.65% (z = 4.4; p < 0.001) and against control 0.165% (z = 5.0; p < 0.001) (Figure 4). The mean sperm-egg ratio for the 3 dilutions treatment were respectively 1.1×10^7^ ± 1.96×10^5^, 1.1×10^6^ ± 1.96×10^4^ and 1.1×10^5^ ± 1965 sperm per egg. The mean sperm concentration in the activation medium (4 mL) was respectively 2.2×10^7^ ± 3.9×10^5^, 2.2×10^6^ ± 3.9×10^4^ and 2.2×10^5^ ± 3,930 sperm per milliliter. The mean (± S.E.) sperm concentration of the males was 2,675 ± 309 Mio/ml.

**Table 2.**
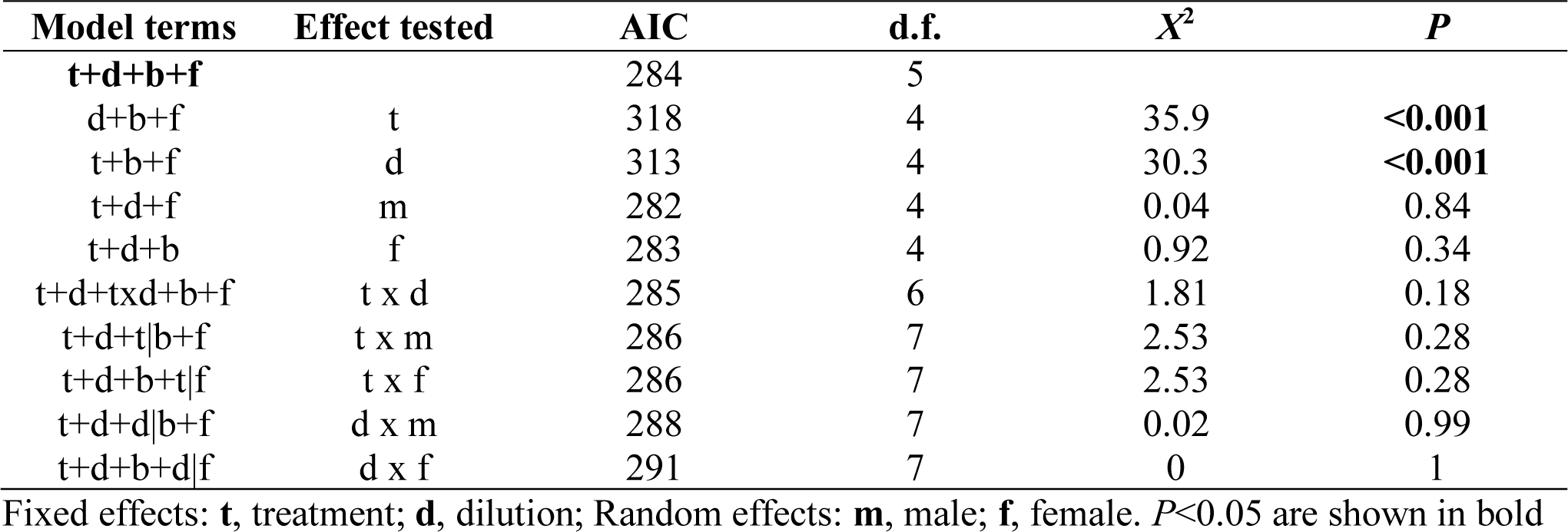
Likelihood ratio tests on mixed-effects model regressions on fertilization success. Models including or lacking the term of interest were compared to reference models in bold to determine the significance of the effect tested.

**Figure 4.**
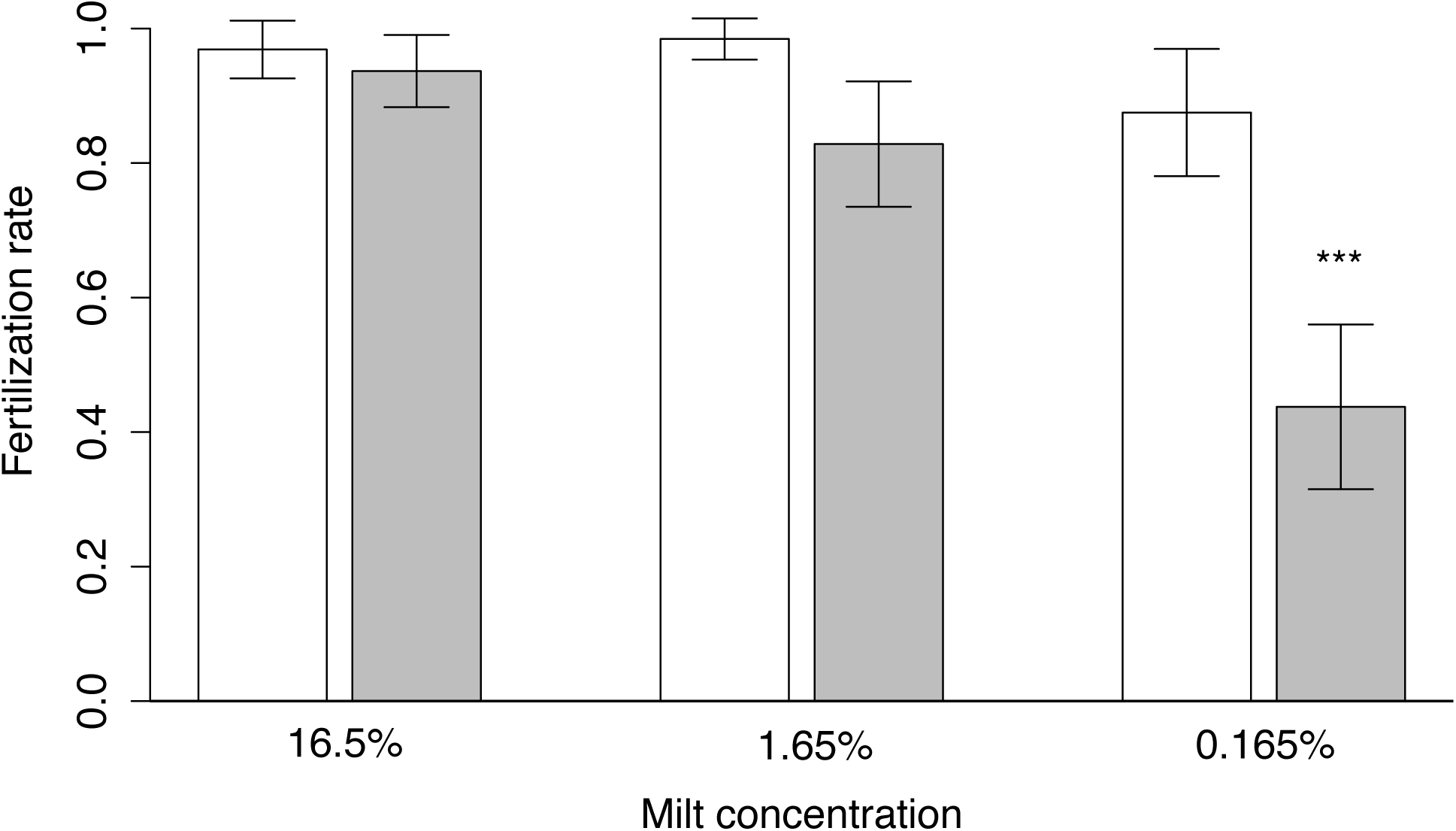
Mean fertilization success in the second experiment. The 3 density treatments are indicated on the x-axis, the white bars indicate fresh sperm and the grey bars frozen-thawed sperm. Error bars indicate 95% confidence interval.

## 4. Discussion

We found that an extender composed of 10% methanol and 0.15M glucose was highly effective in brown trout, leading to fertilization success similar to that of fresh sperm even at high dilution. These findings support previous ones [7,8]. We here compared the effectiveness of this simple extender to a common DMSO-based solution while controlling for parental effects. In the first experiment, we found significant maternal effects on fertilization success. Such effects are typically found in salmonids [14–16] and will not be discussed in the present paper.

As expected from the manufacturer instructions, we reached a fertilization success of about 80% of what is obtained with fresh semen using the commercial DMSO-based extender. Sperm frozen in the methanol-glucose extender performed significantly better and in fact as good as fresh sperm in our first experiment. This confirmed the suitability of methanol-glucose as an effective extender for the cryopreservation of brown trout semen. In practice, this extender showed two main other advantages over the DMSO-based extender.

First, DMSO is toxic and minimizing the time between mixing and freezing is important. However, there are no such time constraints with the methanol-glucose extender. Second, we observed that working with DMSO in a cold environment such as a hatchery (range from 2-10°C) is not easy. DMSO has its fusion point at 18.5°C. Therefore, we had to store and prepare the solutions in a warmer place. None of this problem was encountered with methanol which has a freezing point at -98°C and which was shown to be the least toxic cryoprotectant in loach when tested against DMSO, glycerol, and ethylene glycol [17].

A major problem for cryopreservation in a hatchery is the volume of eggs to be fertilized. Due to the dilution in the extender and the size of a straw, the absolute volume of sperm available per straw is low. There are several ways to overcome this problem. One solution is to increase the size of the straws. The use of 1.2 mL and 5 mL straws was tested before [6][18], leading to satisfying results although fertilization rate remains higher with smaller straws. This is mainly due to the inequality of cooling rate within a straw when its volume increases. Another option is to increase the concentration of the extender in order to change the dilution ratio and increasing the volume of sperm per straw. This was for instance tested by Ciereszko et al. [19] with whitefish semen and methanol-glucose extender. They suggested that a dilution ratio of 3:1 would allow the freezing of more cells per straws although they observed some changes in the motility parameters.

Our approach is that, although sperm egg ratio is diminished, the volume of the fertilization fluid can be increased by diluting the semen after thawing. In our case, we diluted semen after thawing 100-fold, leading to a final concentration of sperm in the fertilization fluid of 0.165%. Our results demonstrate that it is only at the least concentrated dilution that frozen-thawed sperm showed diminished fertilization ability. At this dilution, the sperm egg ratio was of 110,000:1. The lowest sperm egg ratio with frozen-thawed sperm not reducing fertilization success that we found reported in the literature is of 300,000:1 for brown trout [8,20]. However, the amount of eggs per clutch (n = 8) used in our study was low compared to other studies (typically around 200). Although the sperm egg ratio is strictly influenced by the number of both spermatozoa and eggs, the volume of fluid at the moment of fertilization may also play a role. For a given amount of eggs and spermatozoa, the larger is the volume of the fertilizing solution, the lower is the chance for a sperm to encounter an egg although the sperm egg ratio remains constant. This raises the need of standardization when it comes to the development of protocol, as suggested by Tiersch et al., (2011) [21].

To conclude, the methanol-glucose based extender is more efficient than a common DMSO-based extender for the cryopreservation of brown trout semen if experimentally tested in direct comparison, i.e. controlling for potentially confounding factors. The effectiveness of methanol-glucose based extender allows working with comparatively high dilutions while still reaching the fertilization success that can be expected with unfrozen semen.

## 5. Acknowledgments

We thank C. Küng, B. Bracher and U. Gutmann for permission, access to the fish and facilities as well as I. Castro, D. Zeugin, D. Maitre and A. Uppal for help in the field. The project was funded by the Swiss National Science Foundation (31003A_159579).

